# A mitochondria-targeted antioxidant affects the carotenoid-based plumage of red crossbills

**DOI:** 10.1101/839670

**Authors:** Alejandro Cantarero, Rafael Mateo, Pablo Camarero, Daniel Alonso, Blanca Fernandez-Eslava, Carlos Alonso-Alvarez

## Abstract

The mechanisms involved in the production of red carotenoid-based ornaments in vertebrates are still poorly understood. Those colours generated by red carotenoids often depend on the enzymatic production (ketolation) of these pigments from dietary yellow carotenoids. Recently, it has been proposed that this conversion takes place at the inner mitochondrial membrane (IMM). This implies that carotenoid ketolation and cell respiration could share the same biochemical pathways. Such a link would favour the evolution of red ketocarotenoid-based ornaments as reliable indices of individual quality under a sexual selection scenario. We exposed captive male red crossbills (*Loxia curvirostra* Linnaeus) to two different synthetic antioxidants designed to penetrate into the IMM: a synthetic ubiquinone (mitoQ) and a superoxide dismutase mimetic (mitoTEMPO). MitoQ decreased the blood levels of substrate yellow carotenoids and tocopherol. This could be attributed to the characteristics of the mitoQ molecule, which can distort the IMM structure, increasing free radical (superoxide) production and, potentially, antioxidant consumption. Contrarily, mitoTEMPO-treated birds increased the plasma levels of the second most abundant red ketocarotenoid of red crossbills (i.e. canthaxanthin). MitoTEMPO also increased plumage redness and total ketocarotenoid concentration in feathers among those birds exhibiting a redder plumage at the beginning of the study, rising the plasma values of the main red pigment (3-hydroxyechinenone) in paler birds. The results as a whole support the involvement of the mitochondrial antioxidant machinery in carotenoid biotransformation. The fact that the initial plumage redness determined the effect of mitoTEMPO suggests that the mitochondrial-based mechanism is intimately linked to individual quality.

**Summary statement:** Antioxidants designed to penetrate the mitochondrial membrane increased avian plumage redness but depending on pre-existing colouration. This supports mitochondrial involvement in the evolution of carotenoid-based ornaments as reliable quality signals.

## Introduction

How and why animal conspicuous colourations evolve are recurrent questions from Wallace’s aposematism concept (Wallace, 1877) and Darwin’s sexual selection (Darwin, 1871). Among showy colours exhibited by vertebrates, those produced by carotenoid pigments (many yellow-to-red ones) have attracted the most attention due to three particularities of these compounds (1) they cannot be synthesized by the animal organism from other substrates, thus being exclusively obtained with the diet, (2) they are theoretically scarce in food resources, which would make difficult its acquisition, and (3) they have physiological functions contributing to maintain homeostasis (e.g. Britton et al., 2004; von Schantz et al., 1999). This triangle has allowed formulating hypotheses explaining the evolution of coloured traits as reliable signals of individual quality in mate choice/intra-sexual competition contexts (see also Blount et al., 2009 for aposematism). These hypotheses establish trade-offs in the allocation of time/energy to searching for carotenoid-rich food vs. self-maintenance, and in the allocation of ingested carotenoids to colouration vs. homeostasis (see Endler, 1980; Grether et al., 1999; Kodric-Brown, 1985; Lozano, 1994; von Schantz et al., 1999). An inefficient trade-off solution would induce disproportionally higher costs for low-quality animals, making carotenoid-based coloured traits signals of individual quality (Grafen, 1990; Hasson, 1997; see also recently Penn and Számadó, 2019).

Some authors have, however, argued that dietary carotenoids could not be so scarce, at least among avian species (e.g. Hill and Johnson, 2012; McGraw, 2006). We must here note that first support for the scarcity assumption was provided from fish studies (Endler, 1980; Grether et al., 1999; Kodric-Brown and Brown, 1984). That argument would deactivate the trade-off-based hypotheses, removing those costs derived from the investment of resources in colouration (Hadfield and Owens, 2006; Koch and Hill, 2018; Koch et al., 2019; Svensson and Wong, 2011).

Particularly, Geoffrey Hill (2011) postulated that, instead of resource-allocation trade-offs, a physiological tight link between cell respiration efficiency and carotenoid metabolism exists, implying that some carotenoid-based coloured signals can directly transmit the intrinsic quality of the bearer (also Hill and Johnson, 2012). This means that these traits would act as quality indices instead of pure signals (see terminology in Biernaskie et al., 2014; also e.g. Maynard Smith and Harper, 2003). The idea was formulated in a broad context, giving a general hypothesis for the evolution of any sexual signal that was termed the “shared-pathway hypothesis” (Hill, 2011).

Nonetheless, the shared-pathway hypothesis was developed from the particular case of ornaments generated by transforming common dietary yellow carotenoids (i.e. yellow xanthophylls) to red (keto)carotenoids by oxidoreductase enzymes supposedly located somewhere into the cell respiratory chain (Johnson and Hill, 2013). This theoretical framework was generated from some studies in house finches (*Haemorhous mexicanus*). In that species, males exhibit yellow to red plumages produced by xanthophylls (such as β-cryptoxanthin) or ketocarotenoids (mostly 3-hydroxyechinenone; “3HOE”), respectively (Hill et al., 2002; Inouye et al., 2001; Johnson and Hill, 2013). The question was, however, firstly addressed by Otto Völker in 1957. Ornithologists at that time tried to understand why some birds (red crossbills *Loxia curvirostra* in particular) change their red plumage to a yellowish one under captivity conditions (also Weber, 1961). Although the lack of suitable carotenoid substrates in food under captivity could not be fully discarded (see Hudon 1994 and Hill 1994’s discussions), the ornithologists suspected that other factors were implicated. Thus, Völker (1957) proposed that captive birds were unable to correctly perform the redox transformations converting yellow xanthophylls to red ketocarotenoids (β-cryptoxanthin to 3-HOE, such as in house finches) due to flying restrictions under captivity. The idea was virtually overlooked during the following decades, probably as apparently unrelated to any evolutionary hypothesis.

Currently, studies supporting the shared-pathway hypothesis have begun to appear. A gene coding for an oxidoreductase (ketolase) enzyme candidate for yellow to red carotenoid transformations has recently been described in birds (Lopes et al., 2016; Mundy et al., 2016). Moreover, high levels of yellow and, particularly, red carotenoids have been found in the hepatocyte inner mitochondrial membrane (IMM) of house finches (Ge et al., 2015; Hill et al., 2019). The liver seems to be the main carotenoid transformation site in house finches, red crossbills and other Carduelinae species (del Val et al., 2009b and cites therein; Hill and Johnson, 2012). The accumulation of red carotenoids into the mitochondria supports the shared pathway hypothesis as this organelle is responsible for cell respiration (Hill 2011), and particularly supports the “inner mitochondrial membrane carotenoid oxidation hypothesis” (IMMCOH) as a particular case of the main hypothesis (Johnson and Hill, 2013).

In the IMMCOH, red ketocarotenoids are considered molecularly similar to ubiquinone, which is a key molecule in cell respiration transferring electrons across the respiratory chain. Thus, transforming yellow to red pigments would share the machinery involved in ubiquinone biosynthesis and/or ubiquinone:ubiquinol redox cycle and, hence, affect respiratory pathways (Johnson and Hill, 2013). An experimental finding in another classical model, the zebra finch (*Taeniopygia guttata;* Vieillot 1817), has provided additional support to the IMMCOH. Males bear a conspicuous red bill mostly coloured by canthaxanthin and astaxanthin ketocarotenoids (McGraw and Toomey, 2010). Cantarero and Alonso-Alvarez (2017) administered a synthetic compound targeted to mitochondria to captive males. This substance (mitoquinone mesylate or mitoQ; Smith et al., 2003) introduces the redox-active aromatic ring of ubiquinone (i.e. benzoquinone) into the IMM (e.g. Murphy and Smith, 2007, see Fig. 1). Zebra finches treated with mitoQ improved bill redness (Cantarero and Alonso-Alvarez, 2017).

**Fig. 1.**
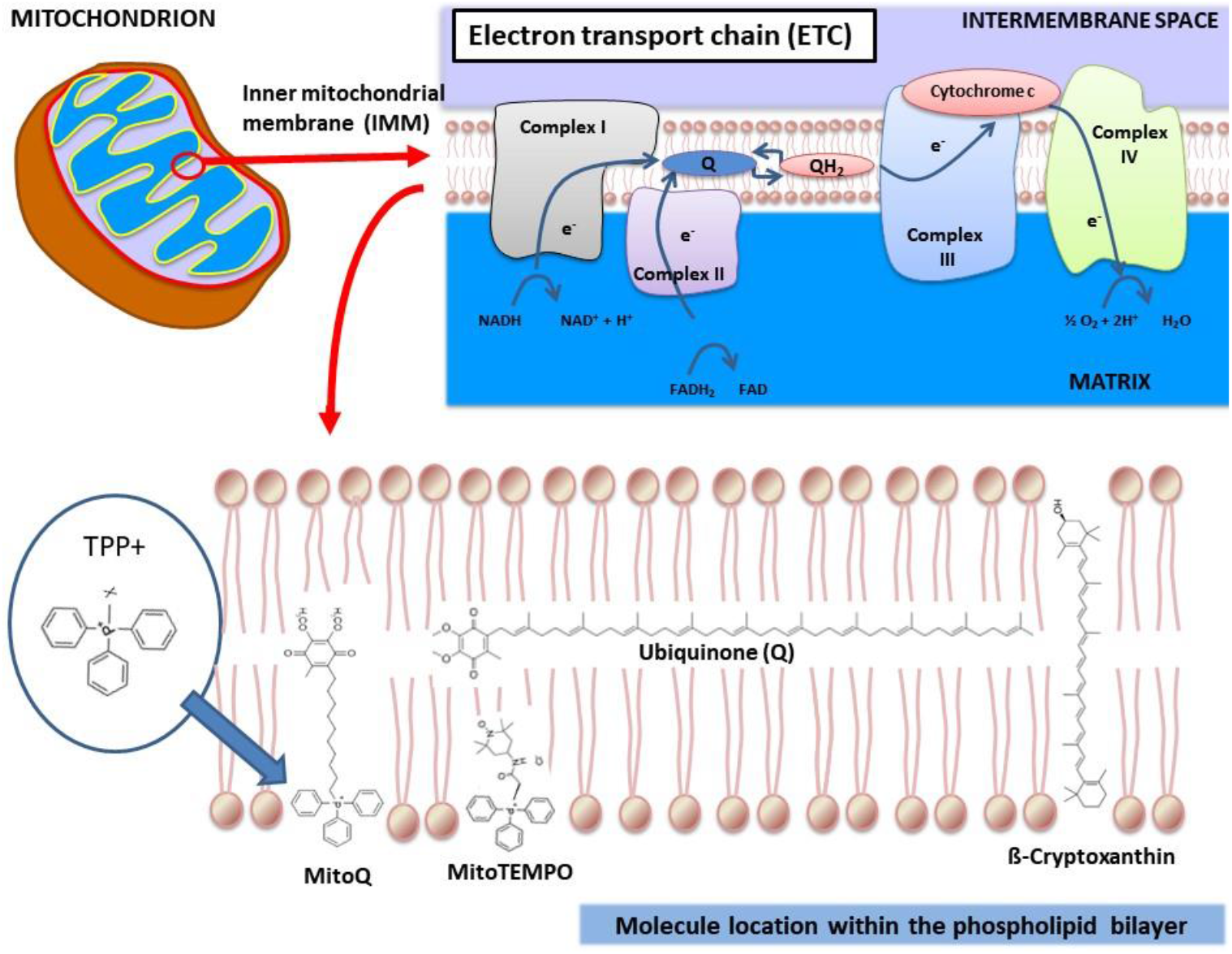
Molecular structure of the two mito-targeted antioxidants and their situation into the inner mitochondrial membrane and electron transport chain.

Synthetic ubiquinone introduction into the IMM is attained by joining together the antioxidant benzoquinone with a triphenylphosphonium cation (TPP^+^) that favours molecule penetration into the IMM and its accumulation in the matrix side (Murphy and Smith, 2007). This procedure has not only been applied to mitoQ production but to many other mito-targeted compounds (Murphy and Smith, 2007; Zielonka et al., 2017). In mitoQ, the active benzoquinone is connected to TPP^+^ by a linker group formed by a 10-carbon alkyl chain (i.e. decyl-TPP^+^ or dTPP^+^). The length of this molecule, however, seems to be responsible for increased membrane permeability, inhibition of the electron transport chain (ETC) and higher superoxide radical generation (see Gottwald et al., 2018; Reily et al., 2013; Trnka et al., 2015). Interestingly, zebra finches only treated with dTPP+ developed paler bills than controls (Cantarero and Alonso-Alvarez, 2017). This opens the question about how strong should have been the ubiquinone effect on colouration without the interfering role of the alkyl chain.

Here, we tested the impact of two mito-targeted antioxidants on the plumage and tissue (blood and feathers) levels of carotenoids and antioxidant vitamins in male birds from the same species used by Völker (1957) and Weber (1961) in their seminal studies: the red crossbill. Although mitochondrial activity was not assessed, the level of specific substrate and product (transformed) carotenoid pigments in blood and feathers allowed us to infer the activity of biotransformation enzymes supposedly located at the IMM. This replacement procedure avoided sacrificing wild birds as a biopsy would have been needed to extract the liver mitochondria. Moreover, as an alternative to mitoQ, mitoTEMPO (2-(2,2,6,6-Tetramethylpiperidin-1-oxyl-4-ylamino)-2-oxoethyl)triphenylphosphonium chloride; Dikalova et al., 2010) was also tested. This molecule includes a piperidine nitroxide that recycles ubiquinol (the reduced ubiquinone form) to ubiquinone (see Fig. 1; Trnka et al., 2008). This action reduces oxidative stress, decreasing superoxide radical levels in mitochondria (Dikalova et al., 2010). In contrast to mitoQ, the nitroxide is joined to TPP^+^ by a shorter linker group than mitoQ, theoretically inducing a lower distortion of the IMM structure (Fig. 1).

We predicted that mitoQ-treated birds, but particularly, mitoTEMPO-treated ones, should develop a redder plumage and higher red ketocarotenoid concentrations in both blood plasma and feathers than controls. Moreover, taking into account the signalling theory (e.g. Biernaskie et al., 2014; Maynard Smith and Harper, 2003), we predict that the intrinsic individual quality should affect the efficiency of the treatments. To address this, the sample was divided by plumage redness before the experiment, categorizing high- or low-redness crossbills that should theoretically represent high- or low-quality individuals (e.g. Galván et al., 2015 for a similar approach). If the availability of substrate carotenoids is unconstrained in food, the addition of mitochondrial antioxidants should improve colouration. We may *a priori* predict that mito-targeted antioxidants should only or mostly benefit low-quality individuals as they could, in some way, be constrained in colour expression. Low-quality individuals could not only be constrained in mitochondrial mechanisms linked to carotenoid biotransformation but in other processes such as the capacity to absorb substrate carotenoids or transport them in the blood (e.g. Madonia et al., 2017). In that case, circulating levels of β-cryptoxanthin or other potential substrates (e.g. β-carotene; see McGraw, 2006, p. 215) should be lower in low-redness birds compared to high-redness ones. We must note that yellow carotenoid uptake is not only a passive diffusion mechanism but also depends on specific receptors in enterocytes (Hill and Johnson, 2012; Jlali et al., 2014; Toomey et al., 2017).

## MATERIAL AND METHODS

### Experimental design

Male red crossbills were captured by means of mist nets during March 2017 at three different locations (i.e. Isaba, Leire and Lakuaga) at the Pyrenean region (Northern Navarra; Spain). The age was determined at the field by the moult pattern (Jenni and Winkler, 1994). Juveniles were excluded from the study. Forty-seven males were finally used. The sample included eighteen 1st year autumn/2nd year spring (8-12-month-old) males (38%). The forty-seven birds were transferred to facilities also located at the Pyrenean region on the same day of capture (i.e. Instituto de Formación Agroambiental, IFA; Jaca, Huesca, Spain). All the birds were housed in four contiguous aviaries. These were 5.9 × 2.5 × 2.2 m (length, width, height, respectively; i.e. 14.75 m^2^ surface, 32.45 m^3^ volume each one). Every aviary was partially covered with a roof (4 m^2^) to provide protection during rainy days. Food was composed of *Pinus sylvestris* pine nuts (Vilmorin, France) and cones, as well as sunflower and hemp seeds. Food, water and grit were all provided ad libitum. Pine branches with leaves were added for behavioural enrichment. The experiment started after one month approximately to allow acclimation (mean 33 days, range 30-56 days). The first manipulation (carotenoid supply) started on May 15^th^, 2017. Birds were captured when required by using a shade net that transformed each aviary in a funnel. This minimized capture stress because less than five minutes were needed to capture all the birds.

### Dietary carotenoid supplementation

Red crossbills are specialized conifer seed feeders, but it is unclear what food item (e.g. pine leaves, resin) is the dietary source of the main substrate carotenoid (i.e. β-cryptoxanthin) involved in producing the red pigment 3HOE (del Val et al., 2009a; 2009b). In fact, pine cones contain relatively low β-cryptoxanthin amounts (< 0.1 mg/Kg in del Val et al., 2009b), and we did not find other food items that pine nuts into the crossbill gizzard during annual ringing campaigns (Daniel Alonso, personal communication). To discard a loss of redness due to β-cryptoxanthin scarcity, the birds received a dietary supplement. It was made by mixing two products: (1) a dry extract of mandarin (*Citrus unshiu*) that contained β-cryptoxanthin at 0.17 mg/g (ref. 0677, Supersmart; Luxembourg) and (2) a synthetic β-carotene provided in corn oil that contained 291 mg/g of β-carotene and 7 mg/g of tocopherol (β-carotene 30% FS, DSM Nutritional Products, Switzerland). These figures were verified by HPLC analyses such as reported below. The β-carotene supplement was chosen because this compound is more abundant in crossbill natural food than β-cryptoxanthin (del Val et al., 2009b). Moreover, β-carotene can be transformed into retinol but also enzymatically modified to echinenone by one oxidation and then to canthaxanthin by another oxidation step or, instead, to β-cryptoxanthin by one oxidation and one hydroxylation (McGraw, 2006 p. 215).

The synthetic β-carotene solution was diluted in peanut oil (SIGMA-ALDRICH ref. P2144) at 1:100 volumes to reduce its very high concentration. We here used peanut oil because it has previously been used in avian studies manipulating dietary carotenoids (e.g. Saino et al., 2000) and because it allows dilution without adding substantial carotenoid amounts (0.1 mg/kg in crude oil; Carrín and Carelli, 2010). The mandarin dry extract was then added to that solution (19.3 g per 100 mL; i.e. the highest concentration that we were able to dilute), the result being mixed by a vortex and also sonication (30 s) in cold water. The mixture was maintained refrigerated and away from light, and regularly made to avoid oxidation (Alonso-Alvarez et al., 2004). All the birds received the carotenoid mix by pipetting 150 microliters into the mouth every other day from the cited date until June 18 (36-day period).

A group of five red male crossbills (see colour categories below) was used to verify that the carotenoid supplement indeed increased circulating levels of substrate carotenoids. Accordingly, they received the same oil amount (150 microliters every other day throughout the experiment) but containing peanut oil only and no additional pigment. They were also injected with saline only (see injection treatment below) to allow comparisons with the remaining birds. At least one bird of this small group was present in each aviary. The five birds were, nonetheless, excluded from the analyses testing antioxidant treatment effects.

### The antioxidant treatment

The males receiving the carotenoid supplement were randomly assigned to one of the three antioxidant treatments (control, mitoQ or mitoTEMPO, *n* = 14 per group; *N* = 42). The number of birds older than one year was fully balanced among these groups (*χ*^2^ = 0.00, df = 2, *P* = 1.00). All these birds were subcutaneously injected into the back every other day throughout 20 consecutive days. When measured the day before the first injection, no significant difference among treatments, colour categories or its interaction was detected on the tarsus length, body mass or size-corrected body mass (i.e. by adding tarsus length as a covariate to a normally distributed GENMOD testing body mass; all *P*-values > 0.23; see also Statistical analyses section). Each injection had 130 μL volume of saline plus MitoQ or, alternatively, MitoTEMPO. MitoQ was kindly provided by Prof. Michael P. Murphy, whereas mitoTEMPO was purchased from SIGMA-ALDRICH (ref. SML0737). MitoQ was administered at 1mM (2.27 mg/Kg/day), which is the dose that induced an increase in bill redness in male zebra finches (Cantarero and Alonso-Alvarez, 2017). With regard to mitoTEMPO dosage, we first considered a study where mice reduced mitochondrial superoxide production and oxidative damage in muscles and vascular tissue when they received 1.5 mg/Kg/day in saline subcutaneously injected throughout 12 weeks (Vendrov et al. 2015; see also Nazarewicz et al., 2013 for same dosage and effect as anticancer molecule). We then performed a pilot study involving 10 male zebra finches randomly assigned to five different concentrations (0, 0.334, 0.668, 1.335 and 2.67 mg/Kg/day) subcutaneously injected in saline every two days for three weeks. The highest dose (2.67 mg/Kg/day; 3mM) was finally chosen as we did not find a significant correlation between dose and body mass change (%) suggesting any health impairment (Spearman’s *r* = 0.320, *P* = 0.367), and no evident toxicity symptoms (behaviour changes, fatigue, lack of alertness). Additionally, the change (%) in redness at the upper bill mandible increased with dosage (Spearmans’ *r* = 0.82, *P* = 0.01; see also Cantarero & Alonso-Alvarez 2017 for colour analysis methods).

All the crossbills were weighed and photographed on May 23. The following day a blood sample (150 microliters) was taken from the jugular vein and birds were injected by the first time, that is, ten days after the start of the carotenoid supplementation, which should have allowed birds to circulate enough substrate carotenoids for transformation. Data obtained in these two days (May 23 and 24) provided the initial values of the experiment. Two days after the first blood sampling all the feathers in the rump region were plucked to induce a synchronized feather growth. The injection period ended on June 14^th^, i.e. when every bird had received 11 injections (a three- week exposure, approximately). A second (final) blood sample was taken on June 12th to determine the final levels of plasma carotenoids under antioxidant exposure. Birds were allowed to end the feather regrowth until June 23th, when they were again weighed and photographed, and the new rump feathers were carefully removed for quantifying carotenoids. Feather samples were stored at −80°C. Birds were released in the original location when they were again fully feathered (July 10^th^). Four birds were found died before the end of the experiment without clear signs of illness or body mass loss (one control, two mitoQ- and one mitoTEMPO-birds). Mortality did not differ among treatments (*χ*^2^ = 0.55, df= 3, *P* = 0.76). Additionally, the second blood sample of one mitoTEMPO and one mitoQ-bird, as well as the rump feather sample of one mitoQ-individual, could not be analysed due to problems during the laboratory processing. Finally, one control bird was unable to produce enough rump feathers for analyses. Therefore, the final sample sizes were 37, 36 and 35 for colour, feather and plasma analyses, respectively. No colour category × antioxidant treatment combination included less than five birds, and the sample was always balanced (all *χ*^2^ tests: *P* > 0.20).

### Colour measurements

The red crossbill plumage was photographed (Canon EOS 50) by putting the birds always at the same position and fixed distance from the objective (Canon Macro Lens EF 50 mm). A Kaiser Repro Base (Kaiser Fototechnik, Buchen) including a gridded board and a column to place the camera at the same height was used (distance from the board to the lens: 38 cm). The base was covered with opaque grey cellular polycarbonate sheets placed in a vertical position to cover the four sides of the board but allowing to enter the camera objective and ring flash (Canon Macro Ring Lite MR-14EX) from the top. The sheets were perforated to allow entering the hands of a person that would hold the body of the bird resting on the board surface. All the open surfaces were covered with PVC blackout fabric to reduce the light on the board surface. The bird was placed in a prone position by pulling the bill and legs. A standard grey card (Kodak, NY, USA) was used as a reference, being placed next to the bird’s flank on the board surface and always in the same position. The focus and diaphragm of the camera, as well as the ring flash, were all manually fixed to avoid the interference of automatic functions. Digital photographs were standardized and analysed using the recently developed ‘SpotEgg’ software, an image-processing tool for automatized analysis of avian colouration that solves the need for linearizing the camera’s response to subtle changes in light intensity (Gómez and Liñán-Cembrano, 2017). We have previously shown that picture-based measurements are highly correlated with the redness measurement (i.e., red hue) obtained from portable spectrophotometers (Alonso-Alvarez and Galván, 2011; Mougeot et al., 2007). SpotEgg, however, allows the user to manually draw any region and provide information about its colouration, shape or other features (ESM in Cantarero and Alonso-Alvarez, 2017 for additional details). The measure of a large delimited area, as opposed to portable spectrophotometers that analyse the colour of reduced spots (usually 1-2 mm), makes this tool useful for evolutionary biologists aiming to capture most of the variability among individuals (Gómez and Liñán-Cembrano, 2017). Accordingly, for each animal, the average of red, green, and blue components of the rump surface was calculated. We then determined hue values by means of the Foley & van Dam (1982) algorithm. Repeatability (Lessells and Boag, 1987) calculated on a set of digital photographs measured twice (*n* = 30) was very high (*R* = 0.99, *P* < 0.001).

### Initial colour variability in our sample of birds

Large variability in plumage colour was present in our sample, birds showing body feathers ranging from yellow to red and including intermediate phenotypes such as homogeneous orange or individuals with small patches of red to yellow feathers randomly distributed in the body (patchy birds). These male phenotypes are frequent in this species (del Val et al., 2014). To take into account this in our statistical analyses, the redness value of the rump at the start of the experiment was used to divide the sample by its median. This should theoretically classify birds in high- and low- quality signallers (Galván et al., 2015 for a similar procedure). This led to two blocks (23 high-redness and 24 low-redness birds) well-balanced among treatments (*χ*^2^ = 0.76, df= 2, *P* = 0.683; see also ESM).

The age category (8-12-month-old vs older birds; see above) was balanced between the two colour categories in both controls (Fisher’s two-tailed test *P* = 0.301) and mitoTEMPO-birds (*P* = 0.580). However, a trend to significance was found among mitoQ-birds (*P* = 0.091; *n* = 14). Here, low-redness birds only included an individual older than one year (1/8), whereas this age group showed a higher frequency (4/6) in high-redness birds. Therefore, colour-related effects in mitoQ birds could partly have been due to age differences. However, the colour category did not show any significant effect among mitoQ birds (see Results). Moreover, when the two-level age factor was added to the statistical models, final models and conclusions were unaffected (data repository at DIGITAL.CSIC, doi: pending on acceptance).

### Sample processing

Blood was immediately stored in a cold box (1-6°C) and centrifuged within 6 h at 10,000 rcf for 5 min at 4 °C. Plasma samples were then separated and stored at −80°C until the analyses. Carotenoid analysis in plasma was performed as described before in García-de Blas et al. (2011; 2013). Feathers in the regrown rump were carefully plucked and stored at −80 °C. These feathers, when analysed, were defrosted and the yellow-to-red part of each feather was cut by a scissor. The total mass of this feather sample was weighed for each bird (mean 8.4 mg, range 4.4-13.6 mg), and this measure used for concentration calculations. The carotenoids in feather samples were extracted following the method described by McGraw et al. (2003). In order to avoid any external contamination of carotenoids from uropygial secretion, the feathers were washed with ethanol, hexane and then air dried. Then, the feathers were extracted in a mixer mill with grinding steel balls (MM400, Retsch, Haan, Germany) for 3 min with 5 mL of methanol and 50 μl of internal standard (retinyl acetate at 0.5 mM). Then, the extract was filtered through a nylon syringe filter 0.2μm, dried under N_2_ stream and finally re-suspended with 200 μl of ethanol and transferred to a chromatographic vial for HPLC analysis.

### Carotenoid and vitamin quantification

The analyses of carotenoids and vitamins A and E in the rump feathers and plasma were performed by HPLC (Agilent Technologies 1200 series) with DAD and FLD detectors, following the methods described by Rodríguez-Estival et al. (2010), García-de Blas et al. (2011; 2013) and Alonso-Alvarez et al. (2018) with some modifications. Compound separation required a column Zorbax Eclipse XBD-C18 (4.6×150 mm, 5 μm) with a precolumn Zorbax Eclipse XBD-C18 (4.6×10 mm, 5 μm). The injection volume was 20 μL. The chromatographic conditions consisted of gradient elution of two phases (A: formic acid 0.1 % in water; B: formic acid 0.1 % in methanol). The initial conditions were 50 % A and 50 % B, changing to 0 % A and 100 % B in 20 minutes, keep this condition 30 minutes and return to initial conditions in 5 minutes, all time the flow rate was 1 mL/min. Standards of lutein, zeaxanthin, canthaxanthin, astaxanthin, 3HOE and β-cryptoxanthin and β-carotene were purchased from CaroteNature (Lupsingen, Switzerland). Retinyl acetate (used as an internal standard) and standards of retinol and α-tocopherol were provided by Sigma-Aldrich. Concentrations were determined from the calibration curve of their standards and linear calibration adjustments. No esterified carotenoids were detected. Carotenoid and vitamin concentrations in plasma or feathers were expressed as nmol/mL and nmol/g for plasma and feathers, respectively.

### Statistical analyses

First, the difference in circulating carotenoid levels between birds fed or not with the carotenoid supplement was tested by non-parametric Fisher exact tests, Pearsons’ *χ*^2^ or Mann-Whitney U tests. This avoided the problem of unbalanced sample sizes (only five birds used as controls here), lack of normality or heteroscedasticity. We must note that the concentration of red ketocarotenoids in plasma showed a high frequency of zero-values, ranging from 36.5% in plasma 3-HOE to 70.7 % in plasma astaxanthin.

Subsequently, generalized linear models were used to test the antioxidant treatment effect and its interaction with the two colour categories (*N* = 35-37, see above). This allowed specifying the distribution type. The GENMOD procedure in SAS 9.4 was used. In these models, sample sizes and variances were well balanced among groups. However, the lack of normality remained due to zero inflation in most ketocarotenoids as well as β-cryptoxanthin and an unknown derivative of β-cryptoxanthin (“UD-β-cryptoxanthin”; see below) both in feathers only. In all these variables, the adjustment to zero-inflated Poisson or, instead, negative binomial distributions were tested (e.g. Ridout et al., 1998). Only the zero-inflated negative binomial (ZINB in SAS) distribution met the no over-dispersion criterion tested by Scaled Pearson’s *χ*^2^. Yellow xanthophylls and vitamins (α- and γ-tocopherol fractions were summed to obtain total tocopherol) in plasma, as well as rump redness, were all normally distributed (a normal distribution specified in GENMOD). Nonetheless, the sum of the five unknown plasma xanthophylls (UX-xanthophylls; see below), as well as the sum of all red ketocarotenoids in feathers, required a log-transformation. Feather astaxanthin was only found in two birds, its variability being not analysed, whereas feather canthaxanthin required a square root-transformation. Finally, the body mass change (%) from the start of the antioxidant treatment to the second blood sampling or, instead, to the date of colour measurement, were normally distributed and also tested by GENMOD for coherence.

The use of the GENMOD procedure prevented to include the aviary as a random factor. However, the three treatment groups (*χ*^2^_6_ = 2.56, *P* = 0.861), the two colour categories (*χ*^2^_3_ = 0.46, *P* = 0.927), or the six groups resulting from its interaction (*χ*^2^_15_ = 0.46, *P* = 0.902) were well-balanced among the aviaries.

With regard to covariates, the carotenoid level at the first blood sampling was tested in GENMOD models analysing carotenoid concentration variability. In the case of the rump colour, the initial redness (before plucking) was not added as a covariate because the colour category captured its variability. Instead, saturation (purity) and brightness (lightness) were included as covariates to assess colour variability fully independent of these parameters (Fitze and Richner, 2002). In any event, the results did not vary when these two terms were removed (treatment × colour category: *P* = 0.025). Initial body mass was also excluded as a covariate in models testing body mass change. Similar results were, anyway, obtained when testing final body mass controlled for an initial weight covariate, but we considered that body mass change provides more useful information. Body mass change models included tarsus length as a covariate to correct for body size variability.

Backward stepwise procedures were used to remove terms at *P* > 0.10. Only two exceptions were made in the models shown in tables: (1) the brightness covariate that was maintained in the redness model to assess hue independently of lightness (see above) and (2) the interaction treatment × colour in the model testing total red carotenoids in feathers as it allowed to check variability among the six group combinations and then visually compare them to that found in the rump redness model (Fig. 5A and B). Satterthwaite-corrected DFs and least-square means ± SEs are reported. LSD tests were used for pair-wise comparisons.

## RESULTS

### Description of carotenoids and vitamins in blood and feathers

In plasma, β-cryptoxanthin, lutein, astaxanthin (but only at the second sampling), canthaxanthin, 3HOE and echinenone were detected. Five unidentified carotenoids were also found. Maximum wavelengths and retention times of these unidentified compounds (see Table S1 in ESM) suggested that they were closer to yellow xanthophylls such as lutein or β-cryptoxanthin. Retinol and α- and γ-tocopherol were also detected. Finally, in the rump feathers, β cryptoxanthin, an unidentified derivative (probably an isomer) of β-cryptoxanthin (“UD-β-cryptoxanthin”; see ESM), lutein, astaxanthin, canthaxanthin, 3HOE and echinenone were detected. Neither retinol or tocopherol was found in feathers (see also Table S2 in ESM for concentration descriptions).

### Supplementary carotenoid effects

In terms of plasma values, β-cryptoxanthin differed between both groups at both the first and second sampling events (*U* = 0.01, *P* = 0.001 and *U* = 8.00 *P* = 0.001, respectively; other pigments *P* > 0.12). Carotenoid-supplied birds showed β-cryptoxanthin values several times higher than controls at both the first (x12 times: median, range: 0.34, 0.097-0.784 vs. 0.028, 0-0.075 nmol/mL, respectively) and second sampling (x6 times: 0.47, 0-1.063 vs. 0.075, 0-0.312 nmol/mL). No other plasma carotenoid differed (all *P* > 123). Additionally, retinol (*U* = 41.0, *P* = 0.045) and total tocopherol (*U* = 33.0, *P* = 0.018) differed at the second blood sampling, with those birds receiving the supplement showing higher and lower levels than controls, respectively (retinol: 5.74, range 3.13-9.98 vs. 4.43, range 3.98-6.12 nmol/mL; tocopherol: 180.48, range 0-383.6 vs. 241.97, range 179.2-330.5 μM).

In terms of feather composition, no control bird for the dietary supplement (those only receiving peanut oil) deposited β-cryptoxanthin or echinenone in the rump (both absence/occurrence Pearsons’ *χ*^2^-tests showed *P* < 0.01). The other pigments were found in both groups. When variability in concentrations was assessed, the effect on β-cryptoxanthin and echinenone was also significant (Mann-Whitney *U* = 10, *P* = 0.001 and *U* = 30, *P* = 0.013, respectively). Among other feather pigments, only canthaxanthin showed a trend to significance (*U* = 44, *P* = 0.069), with supplement birds showing higher values (mean, range: 4.82, 0-17 vs. 2.12, 0-6 nmol/g, respectively). Finally, the redness of the moulted rump differed as expected (*U* = 21, *P* = 0.005; supplemented birds: 2.36, 0.75-5.56; controls: 0.87, 0-1.78).

No bird in this study attained an intense red rump, even when supplied with high levels of carotenoids in food (see ESM). The colour of the moulted rump declined 35.9% in the whole of the sample, ranging from yellow to pale orange. In fact, the percentage of red pigments in feathers only reached 22% approximately of the total amount of carotenoids (Table S2 in ESM).

### Mito-targeted antioxidant effects on body mass

The birds involved in the antioxidant experiment gained body mass from captivity to the first antioxidant injection (mean ± SE: 3.32 ± 0.82%). From that date, the treatment affected body mass change to the second (last) blood sampling (*χ*^2^ = 70.7, df = 2, *P* = 0.020; tarsus length covariate: *χ*^2^ = 2.62, df = 1, *P* = 0.106). MitoQ-birds gained less mass than controls or mitoTEMPO- birds (both *P* < 0.017; mitoQ: 0.018 % ± 0.945, control: 3.198 % ± 0.944; mitoTEMPO; 3.237 % ± 0.943; control vs. mitoTEMPO: *P* = 0.977). When the body mass change was tested to the date of colour measurement, the treatment again became significant (*χ*^2^= 6.84, df = 2, *P* = 0.033; tarsus length: *χ*^2^= 6.12, df = 1, *P* = 0.013). However, birds in both treatments gained less mass than controls, although pairwise comparisons did not reach significance (both *P* < 0.070; mitoQ: 2.590 % ± 0.903, control: 5.471 % ± 0.871; mitoTEMPO; 2.734 % ± 0.871; mitoQ vs. mitoTEMPO: *P* = 0.909). The colour category or its interaction with treatment were never significant (all *P-values* > 0.10), being removed from these models.

### Mito-targeted antioxidant effects on blood composition

In the case of yellow xanthophylls potential used as the substrate for ketocarotenoid production (Table 1), β-cryptoxanthin levels were the lowest in mitoQ treated birds (Fig. 2A), whereas mitoTEMPO and control birds did not differ (*P* > 0.123). Similar results were found for lutein, UX and total tocopherol (Figs. 2B-D). The colour group did not influence these models (all *P* > 0.40).

**Table 1.** Generalized linear models testing the impact of mito-targeted antioxidant treatments and their interaction with colour category on the level of circulating pigments and vitamins of captive male crossbills.

| **Dependent variable (Distribution)** | **Estimate** | **SE** | **d.f.s** | **$\chi^2$** | **P** |
| --- | --- | --- | --- | --- | --- |
| *Terms in the model* |  |  |  |  |  |
| **β-cryptoxanthin (Normal)** |  |  |  |  |  |
| *Initial value covariate* | 0.571 | 0.206 | 1 | 7.66 | 0.006 |
| *Antioxidant treatment* |  |  | 2 | 32.47 | <0.001 |
| **Lutein (Normal)** |  |  |  |  |  |
| *Initial value covariate* | 0.527 | 0.088 | 1 | 35.73 | <0.001 |
| *Antioxidant treatment* |  |  | 2 | 9.95 | 0.007 |
| **Unidentified Xanthophylls (Normal)** |  |  |  |  |  |
| *Initial value covariate* | 0.570 | 0.0935 | 1 | 37.10 | <0.001 |
| *Antioxidant treatment* |  |  | 2 | 7.77 | 0.021 |
| **Total tocopherol (Normal)** |  |  |  |  |  |
| *Initial value covariate* | 0.718 | 0.141 | 1 | 25.96 | <0.001 |
| *Antioxidant treatment* |  |  | 2 | 18.88 | <0.001 |
| **Canthaxanthin (Negative Binomial)** |  |  |  |  |  |
| *Initial value covariate* | 0.0002 | 0.0001 | 1 | 4.83 | 0.0279 |
| *Antioxidant treatment* |  |  | 2 | 6.63 | 0.0363 |
| **<u>Zero inflation model</u>** |  |  |  |  |  |
| *Initial value covariate* | -0.002 | 0.001 | 1 | 9.59 | 0.002 |
| **3-HOE (Negative Binomial)** |  |  |  |  |  |
| *Initial value covariate* | 0.002 | 0.001 | 1 | 7.73 | 0.005 |
| *Colour category* |  |  | 1 | 1.69 | 0.194 |
| *Antioxidant treatment* |  |  | 2 | 12.69 | 0.002 |
| *Interaction* |  |  | 2 | 9.75 | 0.008 |
| **<u>Zero inflation model</u>** |  |  |  |  |  |
| *Initial value covariate* | -0.012 | 0.006 | 1 | 4.46 | 0.035 |
| *Antioxidant treatment* |  |  | 2 | 4.86 | 0.088 |
| *MitoQ* | 3.374 | 1.548 | 1 | 4.75 | 0.029 |
| *MitoTEMPO* | 0.480 | 1.053 | 1 | 0.21 | 0.648 |

Note: Type of distribution, the slopes for covariates as well as Likelihood Ratio $\chi^2$ tests and P-values are reported. Negative binomial models show the main outputs for counting and zero-inflated parts of the models.

**Fig. 2.**
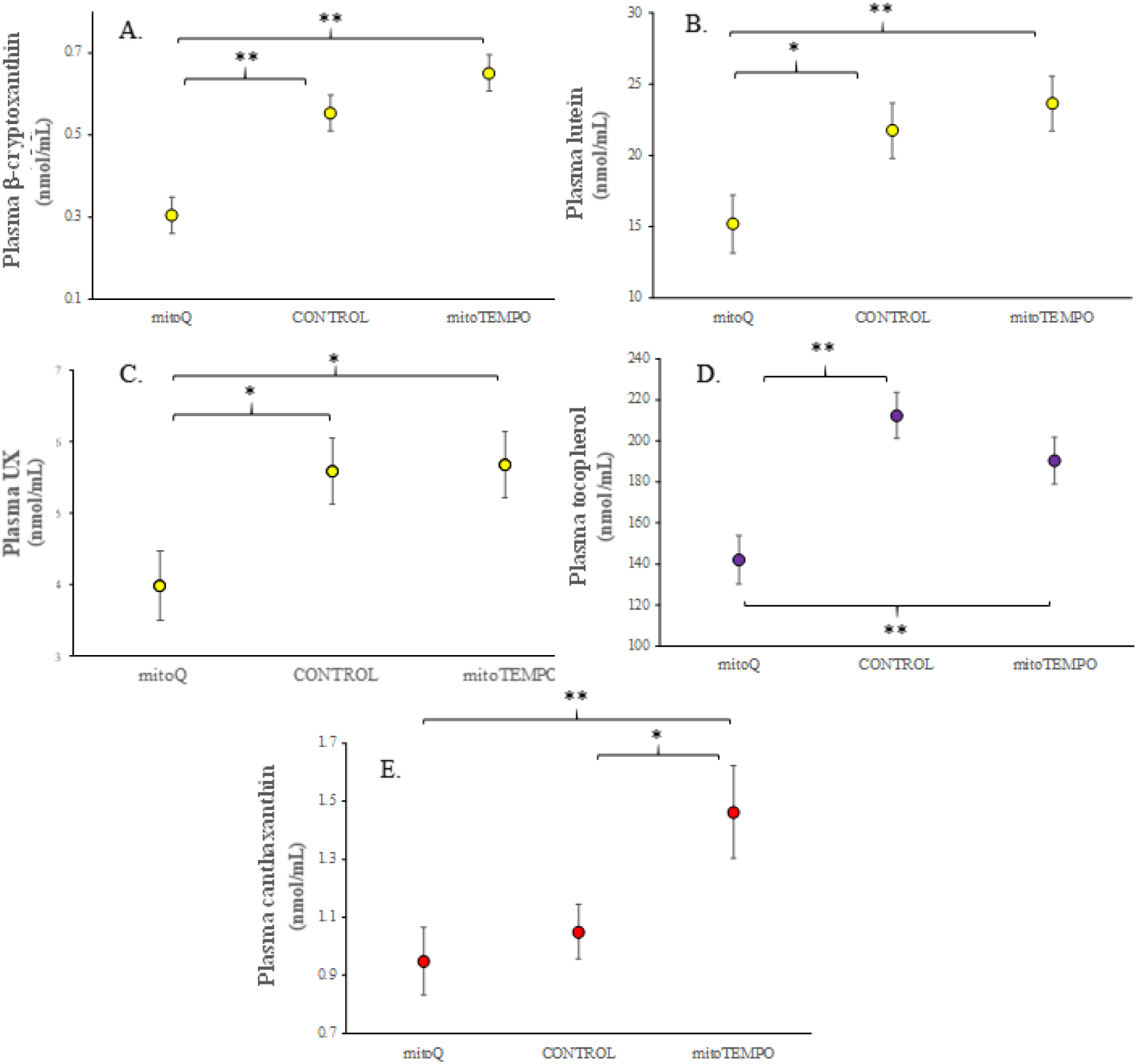
Variability in circulating levels of carotenoids and tocopherol in Eurasian crossbills after treatment with mito-targeted antioxidants. Least square means ± SEs from models. LSD pairwise tests: **: *P* < 0.01, *: *P* > 0.05.

In terms of red ketocarotenoids, mitoTEMPO-crossbills showed higher plasma canthaxanthin values than other groups (Table 1 and Fig. 2E). Only the initial value covariate remained significant in the zero-inflated part of this model as higher initial values led to lower probabilities of zero level at the second sampling (Table 1). The interaction with the colour category was removed at *P* > 0.22.

In contrast, the model testing 3HOE detected a significant interaction (Table 1; Fig. 3). Low-redness mitoTEMPO birds showed significantly higher values than high-redness birds in the same treatment, whereas the same comparison in controls tended to significance in the opposite direction (*P* = 0.093; Fig. 3). Low-redness mitoTEMPO birds also showed significantly higher 3HOE values than low-redness birds from the other two groups and high-redness mitoQ-birds. The latter group showed the lowest mean 3HOE values, and circulated significantly less 3HOE than high-redness controls or high-redness mitoTEMPO- birds (but the latter *P* = 0.057; other comparisons: *P-values* > 0.12). Furthermore, the probability of attaining a zero value was higher in mitoQ-birds compared to controls (*P* = 0.029 in Table 1). To conclude, the other two red pigments in plasma (i.e. astaxanthin and echinenone), as well as retinol, did not show any significant effect (all *P-values* > 0.10).

**Fig. 3.**
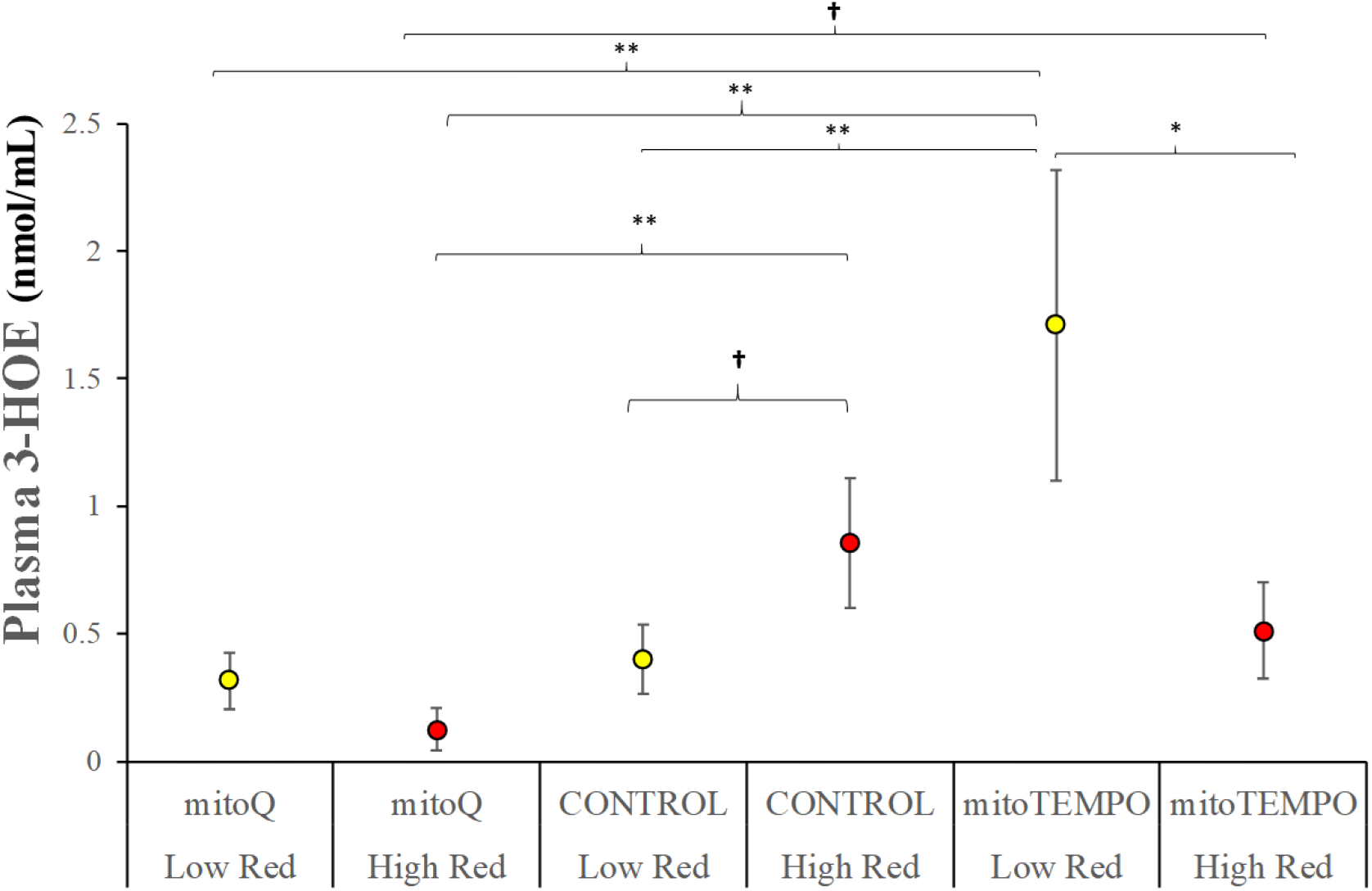
Variability in circulating levels of the red ketocarotenoid 3HOE in red crossbills after treatment with mito-targeted antioxidants. High- or low-redness groups identify those birds over or below the median value of rump redness at the start of the experiment. Least square means ± SEs from models. LSD pairwise tests: **: *P* < 0.01, *: *P* > 0.05; †: *P* < 0.10.

### Mito-targeted antioxidant effects on feather pigments

In rump feathers, the xanthophyll models, i.e. β-cryptoxanthin and UD-β-cryptoxanthin, and the model testing echinenone only retained the treatment factor (Table 2), with mitoQ-birds showing lower levels than the other groups (mitoTEMPO vs control always *P* > 0.10; Figs. 4A-C). The zero-inflated part of these models did not retain any term (all *P’s* > 0.15).

**Table 2.** Generalized linear models testing the impact of mito-targeted antioxidant946 treatments and colour category on the values of feather carotenoids and rump 947 coloration of captive male Eurasian crossbills.

| **Dependent variable (Distribution)** | **Estimate** | **SE** | **d.f.s** | **$\chi^2$** | ***P*** |
| --- | --- | --- | --- | --- | --- |
| *Terms in the model* |  |  |  |  |  |
| **β-cryptoxanthin (Negative Binomial)** |  |  |  |  |  |
| *Antioxidant treatment* |  |  | 2 | 14.92 | 0.001 |
| **UD-β-cryptoxanthin (Negative Binomial)** |  |  |  |  |  |
| *Antioxidant treatment* |  |  | 2 | 10.51 | 0.005 |
| **Echinenone (Negative Binomial)** |  |  |  |  |  |
| *Antioxidant treatment* |  |  | 2 | 12.03 | 0.002 |
| **Log-Total red ketocarotenoids (Normal)** |  |  |  |  |  |
| *Colour category* |  |  | 1 | 0.01 | 0.957 |
| *Antioxidant treatment* |  |  | 2 | 2.21 | 0.331 |
| *Interaction* |  |  | 2 | 3.58 | 0.167 |
| **Rump redness (Normal)** |  |  |  |  |  |
| *Saturation* | 6.409 | 4.790 | 1 | 1.79 | 0.181 |
| *Brightness* | -6.145 | 6.408 | 1 | 3.54 | 0.059 |
| *Colour category* |  |  | 2 | 1.5 | 0.221 |
| *Antioxidant treatment* |  |  | 2 | 3.89 | 0.143 |
| *Interaction* |  |  | 1 | 7.93 | 0.019 |

Note: Type of distribution, the slopes for covariates as well as Likelihood Ratio $\chi^2$ tests and *P*-values are reported. Negative binomial models show the main outputs for counting and zero-inflated parts of the models.

**Fig. 4.**
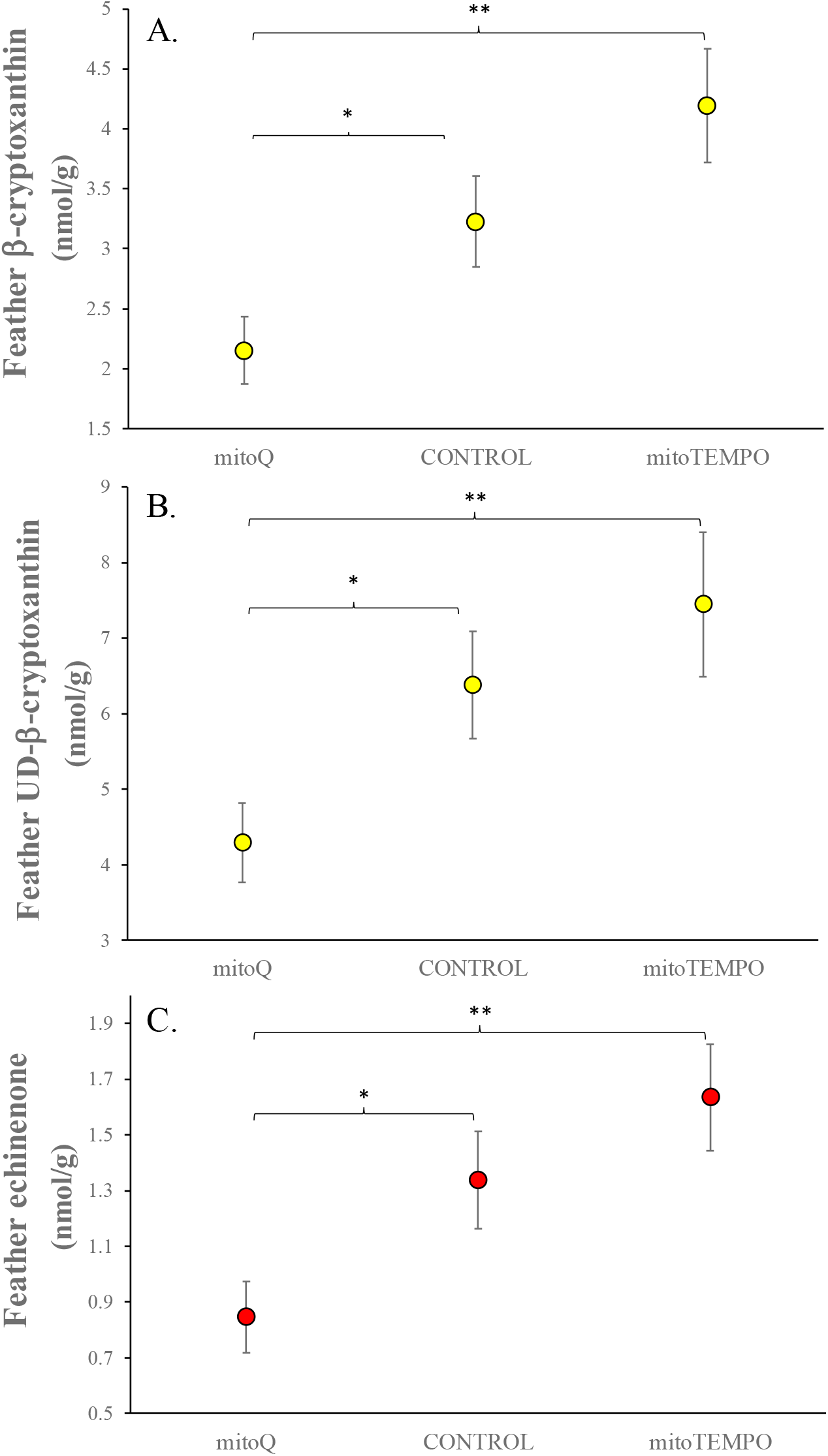
Variability in feather concentration of carotenoids in Eurasian crossbills after treatment with mito-targeted antioxidants. Least square means ± SEs from models. LSD pairwise tests: **: *P* < 0.01, *: *P* > 0.05.

Lutein and Sqrt-transformed canthaxanthin models did not show any significant effect (all *P’s* > 0.12) and are not reported. Furthermore, 3HOE levels were apparently unaffected by the treatment or its interaction with colour category (all *P*-values > 0.40). Only a weak trend to significantly higher levels in high- vs low-redness birds was found (χ^2^ = 2.58, *P* = 0.108; 36.16 ± 8.29 and 22.22 ± 4.41 nmol/g, respectively).

However, if the total level of red pigments in feathers is tested instead, the interaction did not reach significance (Table 2), but mitoTEMPO-birds showed significantly higher values than controls among high-redness individuals (*P* = 0.038; Fig. 5A). The first group also showed a trend to significantly higher values than high-redness mitoQ-birds (*P* = 0.087; other *P’ values* > 0.12). An alternative model testing the sum of yellow carotenoids neither detected a significant interaction (*P* = 0.824), and no term remained in the model (all *P* > 0.20; pairwise comparisons: *P’s* > 0.10).

**Fig. 5.**
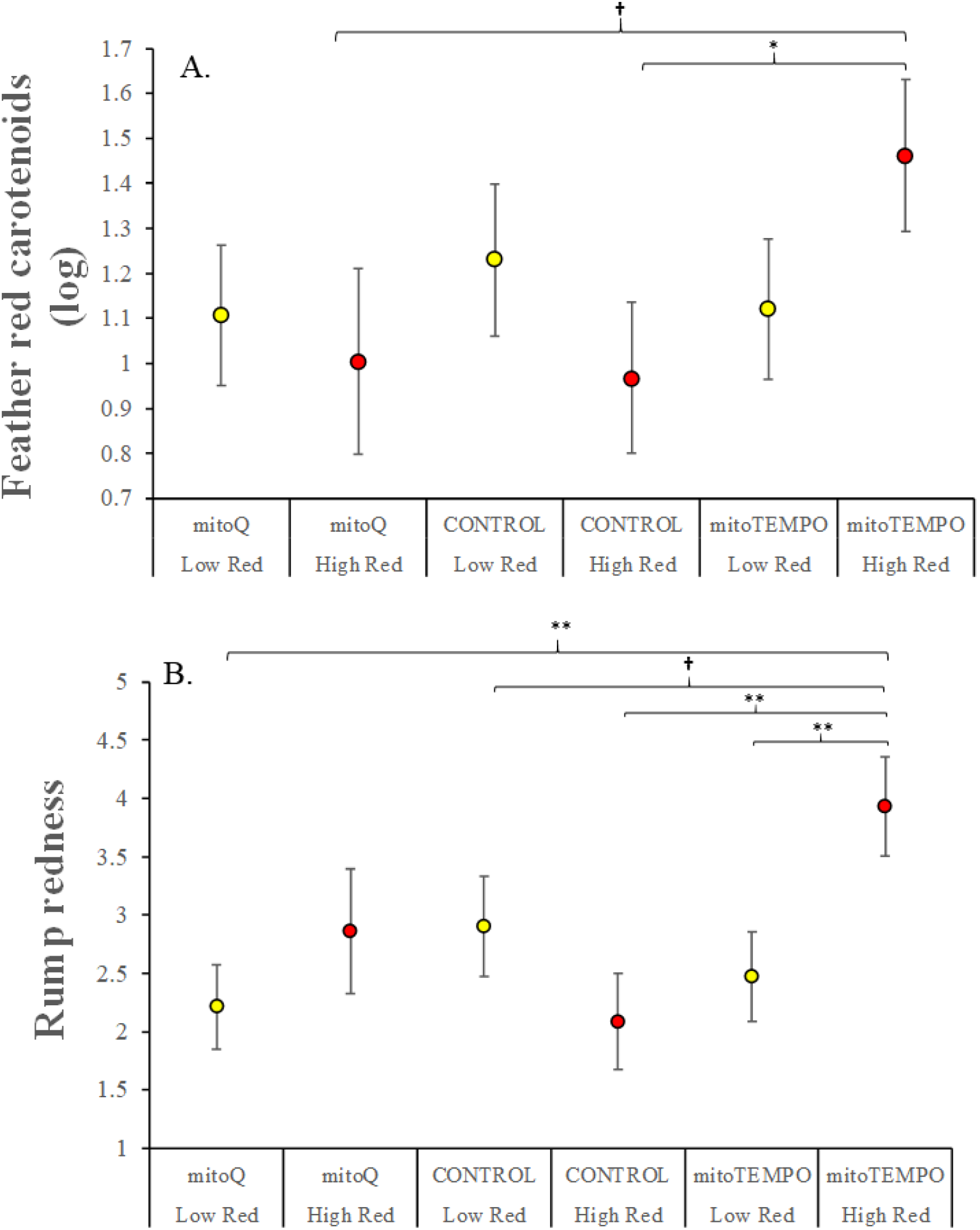
Variability in the total level of red ketocarotenoids in rump feathers (A) as well as rump redness variability (B) in Eurasian crossbills after treatment with two different mito-targeted antioxidants. High- or low-redness groups identify those birds over or below the median value of rump redness at the start of the experiment. Least square means ± SEs from models. LSD pairwise tests: **: *P* < 0.01, *: *P* > 0.05; †: *P* < 0.10.

### Mito-targeted antioxidant effects on rump redness

The interaction between colour and treatment was here significant (Table 2 and Fig. 5B). Agreeing with feather red pigments (Fig. 5A), the rump of high-redness mitoTEMPO-birds was significantly redder than the rump of high-redness controls (Fig. 5B). High-redness mitoTEMPO-birds also showed significantly redder rumps than low-redness mitoTEMPO-birds and mitoQ-birds and a trend to significantly redder rumps than low-redness controls (i.e. *P* = 0.087; other *P’s* > 0.12).

## DISCUSSION

Taking into account the specific characteristics of mito-targeted antioxidants (Dikalova et al., 2010; Murphy and Smith, 2007; Zielonka et al., 2017), our results as a whole seems to support the IMMCOH assumption describing that the mitochondrial membrane is the cell site where yellow carotenoids are biotransformed. Particularly, mitoTEMPO affected rump redness, total ketocarotenoid levels in feathers and canthaxanthin and 3HOE values in plasma. Most of these effects were, moreover, influenced by individual quality assigned by plumage redness before the experiment. However, contrarily to predictions, the treatment improved colouration only among the redder (supposedly high-quality) birds, but not among paler individuals.

Independently of colour categories, we must first note that most of our birds gained body mass probably due to reduced activity and *ad libitum* food. However, birds treated with mito-targeted antioxidants did not gain so much weight as controls. This was first detected in mitoQ-treated crossbills. The same mitoQ-induced effect was found in male zebra finches, although under other housing conditions (i.e. birds paired in cages; Cantarero and Alonso-Alvarez, 2017). Nevertheless, zebra finches only treated with the dTPP+ did not report that effect, suggesting that reduced mass gain is due to changes in redox activity (unpublished results). We can speculate that ubiquinone could increase cell respiration rates, leading to consuming energy stores. In fact, ubiquinone is known to reduce fatness in mammalian models (Allen and Vickers, 2014 and references therein) and mitoQ and mitoTEMPO injections have also shown to reduce body mass in obese rodents (Fink et al., 2017; Gutiérrez-Tenorio et al., 2017; Jeong et al., 2016).

MitoQ-birds also showed lower plasma levels of yellow xanthophylls and tocopherol (Fig. 2). The effect could, at first glance, be attributed to lower food intake. Nonetheless, when body mass change was added as a covariate it always reported *P-values* > 0.25. Tocopherol variability was, however, significantly associated with body mass change, but inversely, decreasing with mass gain (body mass gain covariate: *χ*^2^ = 12.48, *P* = 0.0004; slope ± SE: −6.37 ± 1.80; the antioxidant treatment effect remained significant at *P* < 0.001). We may, alternatively, propose that mitoQ reduced carotenoid and tocopherol intestinal absorption. However, literature in mammals, contrarily, suggests that mito-targeted antioxidants protect the functionality of intestinal cells (e.g. Hu et al., 2018; Wang et al., 2014). As a third option, mitoQ may have increased oxidative stress due to its long alkyl linker group (Fig. 1; Gottwald et al., 2018; Maroz et al., 2009; Reily et al., 2013; Trnka et al., 2015; Yasui et al., 2017). This may have promoted the consumption of tocopherol and xanthophylls to be used as radical scavengers (Britton et al., 2004; Panda and Cherian, 2014). Unfortunately, no measure of oxidative damage was tested, and hence, no firm conclusion can be made here.

The apparently negative (or non-significant) effects exerted by mitoQ on our crossbills contrast with the positive effect of this antioxidant on the bill redness of male zebra finches (i.e. Cantarero and Alonso-Alvarez 2017). This highlights the complexity and diversity of carotenoid-based signalling mechanisms (McGraw 2006). Zebra finches and red crossbills differ in the type of carotenoid-based ornament (bill vs. plumage, respectively) and in the carotenoid biotransformation tissues (bill vs. liver; del Val et al. 2009a,b; Mundy et al. 2016). If we consider that the liver is a vital organ devoted to detoxification and other critical functions, we may hypothesize that a shared (or competitive) pathway involved in sexual signalling (Hill 2011) could have a stronger link to condition when placed at this organ compared to the same pathway functioning on a peripheral tissue (i.e. epidermis). In such case, our mitoQ treatment, even when administered at the same dosage in both species, could have imposed an extra-cost to crossbills that, added to a probable poorer adaptation to captivity (see below), could have prevented to detect a significant effect on bird colour.

In contrast to mitoQ, mitoTEMPO raised the plasma levels of the two most abundant ketocarotenoids of our red crossbills, i.e. 3HOE and canthaxanthin (Table S2). MitoTEMPO-birds should, thus, have been able to increase the carotenoid transformation rate, probably at the mitochondria (Johnson and Hill, 2013). However, the transformation pathway of the two carotenoids differs. Whereas 3HOE values depend on β-cryptoxanthin availability, canthaxanthin would mostly result from β-carotene oxidation (Fig. S1). The absence of β-carotene in the blood of our carotenoid-supplemented birds may support the view that mitoTEMPO increased β-carotene conversion to canthaxanthin. The lack of circulating β-carotene has been reported in several avian species, including passerines fed with the pigment (reviewed in McGraw, 2006). Interestingly, in American flamingos (*Phoenicopterus ruber*), those birds fed with β-carotene did not show circulating β-carotene but increased blood canthaxanthin levels (Fox et al., 1969). We may, therefore, suggest that mitoTEMPO favoured β-carotene transformation at the intestine wall. This would imply the presence of the CYP2J19 monooxygenase (Lopes et al., 2016; Mundy et al., 2016) or some specific β-carotene ketolase probably at the enterocytes. However, such a possibility, as far as we know, has never been reported for any species.

In any event, the most abundant red carotenoid in crossbill feathers is 3HOE (17 v.s 5 nmol/g for 3HOE and canthaxanthin, respectively; see ESM and del Val et al., 2009a). Its main substrate (i.e. β-cryptoxanthin) was well represented in blood and, hence, carotenoid transformation at the liver can, in this case, be defended. Here, low-redness (supposedly low-quality) birds treated with mitoTEMPO showed the highest mean plasma values (Fig. 3), which supports our initial prediction (see introduction). However, rump redness and total ketocarotenoid values in feathers, contrarily, indicate that high-redness birds were the only individuals able to increase colour expression when exposed to the cited mito-targeted antioxidant. In the case of total ketocarotenoid concentrations (Fig. 5A), we must consider the difficulties inherent to carotenoid extraction from feathers and, probably, the low sample size as potential reasons explaining why the colour × treatment interaction was not significant (Table 2). However, high-redness mitoTEMPO-birds showed significantly higher total ketocarotenoid concentrations than high-redness controls, which agrees with feather redness (Fig. 5B).

Our results, intriguingly, suggest that mitoTEMPO did not only promoted carotenoid conversion in low-quality birds (Fig. 3) or in the whole of the sample (see canthaxanthin above) but also ketocarotenoid allocation to feathers among high-quality birds. That differential allocation can be deducted by the fact that, even when showing higher feather redness and ketocarotenoid values than controls, high-redness birds reported lower plasma 3HOE values than low-redness birds when treated with the antioxidant (Fig. 3). Thus, we could argue that the later was due to ketocarotenoids being sequestered by the feather follicles at a higher rate (i.e. a better allocation). In this regard, we performed alternative statistical tests to analyse if circulating 3HOE levels explained (canceled out) the results in feather models. We added plasma 3HOE values at the last sampling or, alternatively, the difference between final and initial 3HOE plasma values, as covariates in the models testing feather redness or ketocarotenoid concentration. These covariates were always significantly and positively correlated to redness and pigment levels in plumage (always *P* < 0.002; slopes ranging from 0.33 ± 0.04 to 0.89 ± 0.11). However, the interaction in the redness model (Table 2) and the comparison between high-redness mitoTEMPO-birds and high-redness controls in the feather ketocarotenoid model remained significant (all *P* < 0.02). This suggests that the cited effects were independent of plasma 3HOE variability, at least at the time of blood sampling. In this regard, we must be cautious when interpreting circulatory values as blood sampling was restricted to a single point (day) after the treatment. Thus, we cannot fully discard that high-redness mitoTEMPO-birds could have previously produced and circulated the highest 3HOE levels favouring feather redness.

We must, anyway, highlight the limitations derived from captivity conditions that could have imposed an extra-cost for all the birds, perhaps making our initial prediction less realistic. In fact, no bird was able to regrowth the intense red rump exhibited by many wild crossbills (Fig. S2 in ESM). This agrees with initial studies in this species demonstrating that captivity severely limits the capacity of these animals to generate red feathers (Völker, 1957; Weber, 1961). In such overstressed conditions, only the best (reddest) individuals would still have been able to show a positive effect when the antioxidant state of the IMM was improved.

In any event, the results as a whole reveal a link between red ketocarotenoid-based colouration and individual quality mediated by mitochondrial antioxidants. Studies simultaneously testing mitochondrial and ketolase activities in model animal species with ketocarotenoid-based ornamentation are now needed to disentangle the mechanism involved in signalling and its ultimate evolutionary consequences. The CYP2J19 discovery (Lopes et al., 2016; Mundy et al., 2016; also Twyman et al., 2018) is exciting. However, although high levels of ketocarotenoids have been detected in the IMM of a similar bird species (Ge et al., 2015; Hill et al., 2019), we need still demonstrate that this or other monooxygenase enzymes in control of carotenoid conversion are indeed placed into the mitochondrial membrane to support the main IMMCOH assumption. Otherwise, if another cell site is involved (e.g. microsomes where the CYP2J family has been described; Nebert et al., 2013), the idea should be rethought to be well placed into the shared-pathway framework (Hill, 2011). Last but not less, our study was not designed to assess if the amount (level) of substrate carotenoids in the body could influence the efficiency of mito-targeted compounds in promoting carotenoid transformations. This is another key point that merits future experimental work if we aim to fully discard a role for resource-allocation trade-offs in the evolution of red ketocarotenoid-based colourations as reliable signals of individual quality.

## ETHICS

This study was approved by the Bioethical Committees of CSIC (Ref. 404/2016) and the local government (Junta de Comunidades de Castilla-La Mancha; Ref. 486728).

## DATA ACCESSIBILITY

All data will be available at DIGITAL.CSIC repository.

## AUTHORS’ CONTRIBUTIONS

C.A.A. and A.C. carried out the experiment, analysed data and drafted the manuscript. C.A.A. designed the study. D.A. and B.F.E. were responsible for crossbill captures and data measurements in the field. P.C., R.M. and A.C. were involved in HPLC laboratory analyses. All authors approve the publication and are accountable for this work.

## Acknowledgments

We are grateful to Prof. Michael P. Murphy for kindly providing MitoQ and advice about its properties. We also thank Dr. Alberto Velando for earlier ideas on the use of mitoQ in testing evolutionary questions. We are grateful to Prof. Geoffrey Hill for advice on dietary carotenoid sources for our captive crossbills. We thank Pilar García Morchón and IFA staff for allowing to use their facilities as well as to Guillermo Mercé Arévalo for regularly obtaining pine-nuts and other aviary material. We finally thank Gustavo Liñán-Cembrano for support with SpotEgg software.

